# The Impact of Habitat Impermanence on Metapopulation Viability and Size

**DOI:** 10.1101/2025.04.09.647889

**Authors:** Emma J Walker, Austin J Phillips, Benjamin Gilbert

**Affiliations:** University of Califoria, Santa Cruz; Woods Hole Oceanographic Institute; University of Toronto

## Abstract

Habitat is often impermanent causing the amount and spatial distribution of habitat patches available to species to vary through time. Theory calls for metrics that fully account for the impact of habitat impermanence on metapopulations, yet all use averages that homogenize spatiotemporal impacts of habitat impermanence in some manner. We develop a novel modelling approach using a deterministic variation of the widely used spatially realistic Levins model paired with a continuous time Markov chain to capture the stochastic impacts of habitat impermanence on finite metapopulations. From this model we derive analytic expectations of metapopulation viability and size by weighting landscape capacities and equilibrium occupancies by the quasi-equilibrium distribution of habitat configurations exhibited by landscapes characterized by a simplest form of habitat impermanence (i.e. random and independent habitat patch loss and gain at constant rates). These provide expectations of metapopulation viability and size in absence of transient metapopulation dynamics. We then show how dispersal and colonization/extinction rates, relative to rates of habitat change, separately impact metapopulation viability and size. Using simulations of our model, we show the measures we propose invariably improve estimation of metapopulation viability and size from earlier estimates, but ultimately overestimate both when species less able to keep pace with rates of change from habitat impermanence in the landscapes they occupy. Our research identifies the importance of fully accounting for spatial habitat configurations through time, and highlights the importance of transient dynamics as rates of habitat impermanence increase.

## Introduction

All habitats are subject to change, causing the spatial distribution of habitat patches for any species to shift through time (Wu and Louks, 1995, Coulson 2021). Though some habitats persist longer than others, many species must persist through frequent disturbance regimes that cause a fraction of habitat patches to be unavailable at any time, a scenario referred to as habitat impermanence. For example, natural variation in rock pools, pitcher plants and other phytotelmata, tree gaps, and other ephemeral resources and shelter, lead to temporal changes in the spatial distributions of populations (Alvarez-Buylla and Garcia-Barrios 1993, Altermatt et al. 2012, Vanshoenwinkel et al. 2008, Rasic and Keyghobadi 2012, Whitney et al. 2016, Mullineaux et al. 2018, Datry et al. 2016, Ferrari et al. 2017, Larned et al. 2010, Roslin and Koivunen 2001, Schenk et al. 2023, Bengtsson 1993). Furthermore, humans increasingly disturb and drive increased natural disturbances (e.g. through climatic extremes) to populations in historically more static habitats (Ramalho and Hobbs 2012, Zeigler and Fagan 2014, Perry and Lee 2019). The need to understand how habitat impermanence interacts with population dynamics to structure population viability and size is increasingly urgent (Van Teefflelen and Opdam 2012, Zeigler and Fagan 2014, Perry and Lee 2019).

Metapopulation models have been a critical development to understanding how populations persist when habitats are patchy (Doak and Mills 1994), yet most empirical studies consider only spatial rather than spatiotemporal habitat structure (Perry and Lee 2019). Spatially explicit models (SEMs) offer well tested threshold estimates and metapopulation size metrics that have become commonplace in conservation planning (Saura and Rubio 2010, Van Teefflelen and Opdam 2012, DeAngelis and Yurek 2017). Meanwhile, metapopulation theory has long discussed the potential importance of habitat impermanence on local and metapopulation dynamics (Levin and Paine 1974). Diverse models, threshold estimates and metapopulation size metrics have been proposed to account for impacts of this habitat impermanence (Levin and Paine 1974, Hanski 1999, Keymer et al. 2000, Hill and Caswell, 2001, Xu et al. 2005, Wilcox et al. 2006, Cornell and Ovaskainen 2008, Dreschler and Johst 2010, Reigada et al. 2015, McVinnish et al. 2016, Smith 2017), and suggest alongside few empirical studies that the impacts of the temporal impermanence of habitat may well outweigh the impacts of spatial connectivity in decreasing metapopulation viability and size (Keymer et al. 2000, DeWoody et al. 2005, Ross et al. 2008, Hodgson et al. 2009, Van Teefflelen and Opdam 2012, Bertassello et al. 2021). Yet, most of this theory remains untested and the few empirical studies that do test model predictions have found the modelling methods and estimates they employ overestimate metapopulation viability and size, but can only postulate hypotheses as to why (Hodgson et al. 2009, Bertasello et al. 2021). Thus, accurately characterizing and understanding the impacts of habitat impermanence on metapopulation viability and size even in the simplest of models has proven challenging and presents a significant barrier to extending empirical research on metapopulations.

A fundamental challenge for theory lies in accounting for temporal dynamics of habitat impermanence as elegantly as current analytic models account for spatial dynamics. Many existing analytic approaches incorporate spatial realism into metapopulation dynamics but inadvertently eliminate aspects of this realism when assessing viability (DeWoody et al. 2005). DeWoody et al. (2005) (and subsequent extensions, Xu et al. 2005 and McVinnish et al. 2016) integrated a fully spatially realistic metapopulation model with a stochastic habitat model.

However, their approach also homogenizes spatiotemporal dynamics by assuming that metapopulation viability is determined by the spatial structure of the full landscape if all possible habitat were present, weighted by the expected (mean) lifetime of each patch. Perhaps unsurprisingly then, these models conclude local patch lifespans are primary in determining metapopulation viability and size in large habitat networks (DeWoody et al. 2005, Xu et al. 2005, McVinnish et al. 2016). While this approach, which we term “spatiotemporal averaging” is better than ignoring temporal dynamics, it nonetheless cannot capture any interactive effects of spatial dynamics with habitat impermanence. More specifically, the specific spatial configurations of habitat through time is lost with this homogenizing approach. As a simple example, consider the three-patch metapopulation displayed in Fig. 1. The overall effect of having each patch absent for some fraction of time leads to several distinct habitat configurations. The non-linear consequences of habitat loss (Walker and Gilbert 2022) imply that the net effect of these configurations will be distinct from a constant spatial structure that suffers an ‘average’ loss of each patch. In extreme cases, even if a landscape with ongoing habitat losses and gains could support a metapopulation on average, specific configurations of habitat could arise even briefly or fairly rarely yet have great impact.

**Figure 1.**
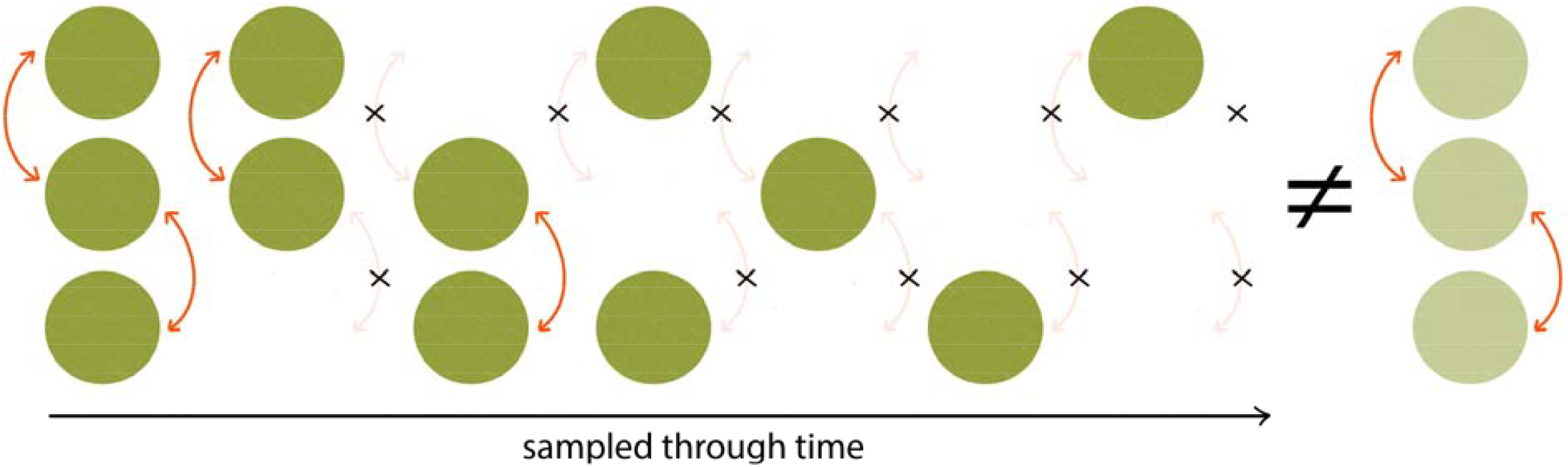
The interactive effects and spatial heterogeneity and habitat impermanence. Current approaches assume that the effects of habitat impermanence can be captured by modeling the spatial configuration of all patches (left-hand side) at their average frequency of occurrence (shaded green; far right-hand side). However, interactive effects of impermanence and spatial heterogeneity can only be assessed if all potential habitat configurations (all dark green scenarios), and their relative frequencies, are jointly considered. In this simplified 3-patch system, each patch can provide colonists to neighbouring patches (arrows), and each patch has an equal probability of being unusable at a given time.

Coupled with the complexity of temporally-shifting habitat structure is the uncertainty of how metapopulations respond to, and average, the effects of extreme events through time. The non-linear responses of metapopulations to habitat loss depends on species traits and habitat configurations (Walker and Gilbert, 2022), meaning the curvature that affects non-linear averaging is dependent both on species and the dynamics of habitat impermanence (Ruel and Ayres, 1999). In addition, the dynamics of habitat loss and gain are necessarily different from a metapopulation perspective. Loss of an occupied habitat patch leads to local extirpation of the populations residing within, whereas creation of new habitat may increase a metapopulation’s expected size but never results in the instant production of an occupied habitat patch. Transient dynamics can cause species to persist long after their metapopulations are no longer viable, (Tilman et al. 1994), and the asymmetric consequences of habitat loss and gain may cause transient dynamics to be more important in the latter scenario. Intuitively, the consequences of transient dynamics will depend on their duration relative to the time spent in individual habitat configurations, but may also depend on whether lags are equally long when metapopulation growth rates are positive or negative. Fully disentangling how these nonlinear and nonequilibrium dynamics alter average metapopulation size is challenging but, as a first step, requires a modelling approach that can be used to determine when they cause metapopulation dynamics to differ from the dynamics expected if transient dynamics were absent.

In this paper, we develop metrics to capture the interactive effects of habitat impermanence and spatial habitat heterogeneity on metapopulation thresholds and size. We use a modelling approach to test our predictions against simulations of average metapopulation size supported in these landscapes subject to the simplest form of habitat impermanence (when habitat patches are randomly and independently gained and lost at constant rates), and contrast with the spatiotemporal averaging approach first developed by DeWoody and colleagues (DeWoody et al. 2005). We first create a model with stochastic habitat dynamics using a Continuous Time Markov Chain (CTMC) governed by rates of patch loss and recovery, meaning that patches are either “lost” and unavailable to support a local population or “recovered” such that they can support a local population. We overlay this habitat model with the dynamics of the metapopulation, modelled using the Spatially Realistic Levins Model (SRLM) (Ovaskainen and Hanski 2001). This choice of a stochastic landscape and metapopulation dynamic model allows us to predict spatiotemporal patterns of habitat configurations and the resulting expected metapopulation dynamics in order to make analytically derived predictions. We then use simulations in which we vary parameters important to spatial and temporal metapopulation dynamics: the rate of patch loss and recovery, the mean dispersal distance of individuals, and the rate at which populations experience local extinction in absence of habitat loss. This allows us to 1) derive improved metapopulation viability and size metrics by weighting these explicitly by the frequency and duration with which particular spatial configurations of habitat arise, as described by the quasi-equilibrium distribution of the habitat’s CTMC, 2) directly relate and compare our estimates of metapopulation size against previous estimates that homogenize spatial-temporal dynamics, and 3) assess when habitat and metapopulation dynamics generate sufficient transient dynamics to alter metapopulation viability and size.

## Model and Methods

The metapopulation model consists of two components. 1) A continuous time Markov chain (CTMC) is used to describe the habitat dynamics. Specifically describing a set of *N* sites suitable of producing viable habitat separated by distances *d*_*ij*_, where habitat is formed with a carrying capacity (*K*) at a rate *r*_*on*_ and lost at a rate *r*_*off*_ (effectively rendering *K* = 0 such that no habitat is available there), 2) A Spatially Realistic Levins Model (SRLM) is used to describe metapopulation dynamics as habitat is formed and lost from the system according to the CTMC. Below, we outline how we use the mathematical properties of the CTMC and SRLM to generate and test the effects of habitat impermanence on metapopulation persistence and size. The model was built and simulated in R (version 4.2.0).

### Habitat Impermanence

Habitat patches can be in one of two states, “on” with carrying capacity (*K*) and supporting local population dynamics, or “off” with no carrying capacity (*K* = 0) or local population dynamics. Different habitat configurations are generated by patches independently turning on and off at rates *r*_*on*_ and *r*_*off*_ where

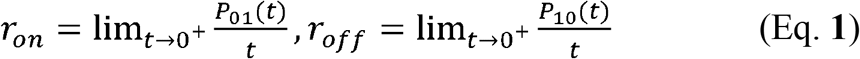

where *P*_10_(*t*) gives the probability a patch is lost at time *t* and *P*_01_ (*t*)gives the probability that it recovers.

The resultant habitat dynamics are described by a continuous time Markov chain (CTMC), with a state space of 2^*N*^ habitat configurations corresponding to every combination of each of the *N* habitat patches being “on” or “off”. For example, considering a maximum of two possible patches, a landscape could be in in any one of four possible configurations: both patches on(*i* = 11), both off (*i* = 00), the first on and second off (*i* = 10), and visa versa (*i* = 01). The change in the probability *P* of being in each possible configuration *j* over time can thus be expressed as

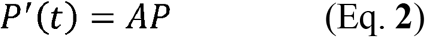

Where *P* is a column vector of the probabilities of being in configuration *j* given a current configuration the landscape *i*. E.g. for a max two patch landscape,

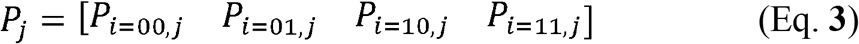

And where A, termed the generator matrix, describes the infinitesimal rate of transition between configurations as individual patches turn on or off. The off-diagonal elements of this matrix *a*_*ij*_, *i* ≠ *j* describe the rates of transition from *i* to *j*, while the diagonal elements *a*_*ij*_, describe the rate of transition out of that configuration given by the negative sum of the transition rates into that configuration (all other elements *a*_*ij*_ in that row). For example, for a system of two patches the generator matrix would be

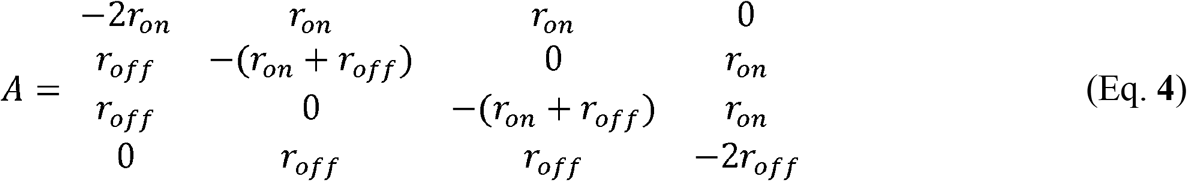

The system of ODE’s described by the Markov chain leads to the solution

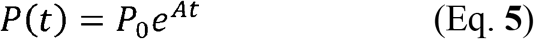

specifying the probability of all configurations through time from an initial state(*P*_0_).

Provided the CTMC process occurs for enough time, it approaches a fixed proportion of time in each configuration and thus samples each configuration with a fixed probability *π* describing the equilibrium of the CTMC, that is

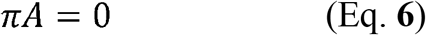

However, species population dynamics cannot occur in total absence of habitat. This means the metapopulation will go extinct instantly when all habitat is lost (if it has not already done so) and the habitat dynamics capable of supporting a metapopulation end when all habitat is lost. Since any later habitat changes occurring are of no consequence to the metapopulation, mathematically, we consider no transition rates out of the ‘no habitat’ state making it an absorbing state to the CTMC. Thus,

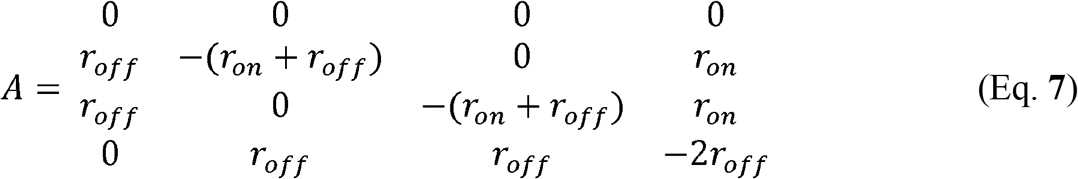

would be a two-patch example of the generator matrix with no habitat as an absorbing state following the general form

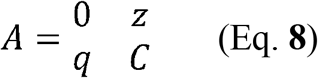

where *z* is a vector of zeros indicating there are no transition rates out of the absorbing state configuration into any of the 2^*N*^−1 other configurations, *q* is a 2^*N*^−1 long vector containing the transition rates into the absorbing state configuration and *C* is a 2^*N*^−1 by 2^*N*^−1 matrix of the transition rates between the non-absorbing (transient) state configurations.

Since there is always a nonzero probability of entering this absorbing state where all habitat is off, the long-term equilibrium of the CTMC is for a species’ habitat itself to eventually reach and stay in this no-habitat state. The time it will take to reach absorption (the no habitat-state) will be longer the more habitat patches are present, and when *r*_*on*_ is large relative to *r*_*off*_. This rate at which the no-habitat state is reached is given by the leading eigenvalue(*ρ*_1_) of *C* and the average time to absorption from each state is the sum of each row of − *C*^−1^ (Darroch and Senata 1967, Kemeney and Snell 1976). However, the time it takes for the habitat CTMC to be absorbed into this no-habitat state, assuming realistic rates of habitat loss and recovery and a reasonably large number of habitat patches, is very long (or neither habitats nor the species they support would persist for very long in nature either).

When time to absorption is long, the habitat CTMC approaches a quasi-equilibrium distribution (QED). The QED, given by the leading eigenvector of *C*, describes the proportion of time the CTMC spends in each of the 2^*N*^−1non-absorbing (transient) configurations of habitat. Since the QED is approached at a rate relative to the absorbing state given by the ratio of the 1st and 2nd largest eigenvaluesρ_2_/ρ_1_ of *C*, it has been shown to generally provide a good measure of the CTMC’s distribution of states when 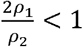 (Artelejo and Lopez-Herrero, 2010). For each combination of *r*_*on*_ and *r*_*off*_ we checked to ensure we are safely within this case throughout the realistic parameter space we consider in this paper 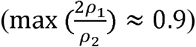.

### Metapopulation Dynamics

The metapopulation dynamics across habitat configurations sampled through time are modelled using Ovaskainen and Hanski’s (2001) Spatially Realistic Metapopulation Model. The SRLM (as formulated in Ovaskainen and Hanski, 2001) describes the stochastic occupancy of habitat patches through time, with patches either occupied or unoccupied at each time-step and the dynamical model specifying the rates at which occupancy changes via colonization and extinction. The SRLM can also be used to model the proportion of habitat occupied through time by using the system of ordinary differential equations (Ovaskainen and Hanski 2003):

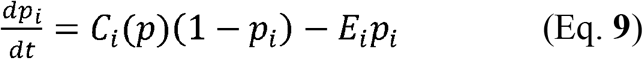

In this deterministic formulation of the model, the metapopulation dynamics (eqn. 9) describe the rate of change in the proportion of habitat occupied (*p*_*j*_) within each habitat patch *i* by the difference at which *i* is colonized versus goes extinct. Here, *C*_*i*_(*p*) is the colonization rate of the unoccupied proportion of patch *j* by dispersers contributed by itself and other patches. This is given by

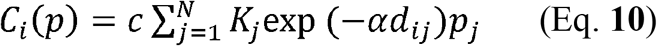

where *c* is the species-specific colonization rate constant, *K*_*j*_ is the carrying capacity of patch *j* that contributes dispersers, *α* is the species-specific parameter inversely proportional to its average dispersal distance, *d*_*ij*_ is the distance between patches *i* and *j*, and *p*_*j*_ is the proportion of patch *j* currently occupied. In the absence of habitat destruction, the proportion of the population lost within a patch is given by the extinction rate,

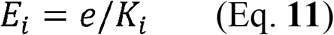

where *e* is the species-specific extinction rate constant and local extinction is inversely proportional to the size of population the patch is capable of supporting *K* as an analytically tractable way of accounting for increased extinction risk of smaller populations due to demographic stochasticity (Hanski 1992).

The dynamics of this model are well studied. An initial vector of patch occupancies *p*_0_ will either tend toward zero (metapopulation extinction) or reach a non-zero equilibrium *p*^*^ (metapopulation persistence) as time increases. Notably, it has been shown that cyclic or chaotic behavior does not occur, irrespective of the number of habitat patches (Ovaskainen and Hanski 2003).

The threshold condition for metapopulation persistence in a landscape (i.e. *p*^*^ > 0) is set by the landscape capacity *λ*_*M*_ which is the dominant eigenvalue of the landscape matrix *M* with elements *m*_*ij*_ which for our model is defined as

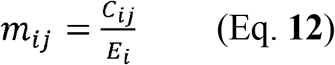

where *C*_*ij*_ is the contribution that patch *j* makes to *C*_*i*_.

In the original SRLM, diagonal elements of *M* were set to zero, meaning that patches could not be recolonized by dispersers from their natal habitat. Given species may also often disperse very short distances such that they never disperse out of or recolonize their natal patch, we model this more biologically realistic scenario of patches to be recolonized also by dispersers of their natal patch by not requiring *j* ≠ *i* in Eqn. (9) and letting 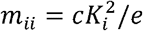 rather than *m*_*ii*_ = 0 (Schnell et al. 2013). Mathematically this change guarantees a real, positive dominant eigenvalue of *M* even when a metapopulation is reduced to one patch for a period of time. The SRLM also requires that the matrix of habitat patches is irreducible, meaning that no patch or group of patches is isolated from all other patches in the matrix. We meet this assumption by assuming that emigrants from each patch can reach all other patches, albeit with extremely low probabilities in some cases.

Defining *δ* = *e*/*c*, the metapopulation persists if *λ*_*M*_ > *δ*, and goes extinct if *λ*_*M*_ < *δ* (Ovaskainen and Hanski, 2001).

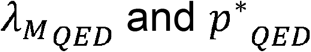

By combining *λ*_*M*_ with the QED, we estimate the full spatiotemporal metapopulation viability.

We derive improved metapopulation viability and size metrics by weighting these explicitly by the frequency and duration with which particular spatial configurations of habitat arise as is described by the QED. Specifically, we calculate the landscape capacity 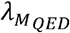and quasi-equilibrium patch occupancy 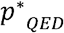 of the stochastic landscape as the geometric mean of *λ*_*M*_ and *p*^*^_*QED*_ (as defined by Ovaskainen and Hanski (2001) for the SRLM) in each static habitat configuration weighted by the quasi-equilibrium distribution of each configuration. Low-density population growth rates are properly captured by their geometric mean (Lande et al. 2003), meaning that the geometric mean of *λ*_*M*_ is appropriate for an SLRM (Ovaskainen and Hanski 2001). Although it is less clear that *p*^*^_*QED*_ should be estimated with the geometric mean, there are a few considerations that lead us to this approach. First, even in highly connected metapopulations, the greater the fluctuations in extinction rates, the more *p*^*^ is expected to fall below the arithmetic mean of *p*^*^ (Levins 1969). Second, the asymmetric role of habitat creation and destruction on extent populations suggests that variation in *p*^*^_*QED*_ will be characterized by growth toward *p*^*^ for each habitat configuration. Finally, should *p*^*^ fall to zero (as expected if *λ*_*M*_ <), it’s long-run expectation will be captured by the geometric mean rather than the arithmetic mean.

To assess the importance of accounting for the full spatial-temporal habitat dynamics, we contrast our metrics with the *λ*_*M*_ and *p*^*^metrics of DeWoody et al. (2005), which in turn has been shown to perform better than other methods (DeWoody et al. 2005, McVinnish et al. 2016). To avoid confusion we refer to *λ*_*M*_ and *p*^*^ calculated using the methods outlined in DeWoody et al. (2005) as 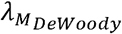 and *p*^*^_*DEWoody*_respectively, and we refer to our metrics as 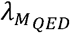 and *p*^*^_*QED*_. Specifically, to do this we have refrained from pulling the scalar out of the landscape matrix *M* in the calculation of *λ*_*M*_ in the calculation of 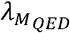 and 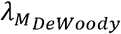. This allows conversion of *λ*_*M*_ to 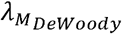 and *p*^*^to *p*^*^_*DEWoody*_by simply replacing the parameter in these calculations with how it is impacted by the landscape dynamics as solved for by DeWoody et al. (2005) as

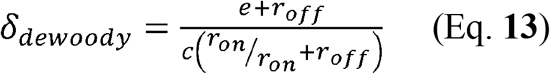

Where now 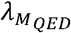 and 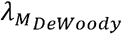 share a common threshold condition for persistence requiring them to be > 1.

### Simulating Habitat Impermanence and Metapopulation Dynamics

We first simulate habitat configurations described by the CTMC, and then numerically solve metapopulation dynamics within and across habitat configurations. We use a Gillespie algorithm to model habitat changes, meaning that we calculate the waiting time ***τ*** between habitat changes.

These waiting times are exponentially distributed with mean *μ* = −*a*_*ii*_ where *a*_*ii*_ are the diagonal elements of the generator matrix A. For simplicity, we refer to these waiting times in units of “years” but note that they could be scaled to match a particular habitat and metapopulation with faster or slower dynamics.

Since local population dynamics require a local carrying capacity(*Kj*) the metapopulation dynamics only occur in patches made available by the CTMC. The initial occupancy of the SRLM in each new configuration of habitat is set with the habitat occupancy it last held in the previous configuration (all patches remaining “on” retain the ***p***_*i*_ held prior to the habitat change, while all newly “off” patches are lost along with the proportion of the metapopulation they supported *K*_*i*_ = 0 so *p*_*i*_ = 0, and all newly “on” patches begin empty *p*_*i*_ = 0). The habitat CTMC models each and every instantaneous change to the habitat meaning only a single patch may blink into or out of existence in an instant but the time *τ* between these instances may be very small. We numerically solved the change in metapopulation proportion within each patch (eqn. 9) using an ODE solver over the period ***τ*** spent in a habitat configuration before a change.

We simulated metapopulation dynamics until each simulation reached a no-habitat configuration (if that occurred within 10,000 years) or up to 10,000 years. We performed 200 replicates, varying rates of habitat disturbance (*r*_*off*_)ranging from1×10^−5^ to 0.1and rates of habitat recovery (*r*_*on*_) ranging from 0.01 to 0.1. This could be considered representative of systems where local disturbances occur on the order of once every decade to once every 100,000 years on average, with habitat recovery occurring on the order of yearly to centennial bases impact annually reproductive species (e.g. for species dependent on hydrothermal venting activity in regions differing by geologic activity or tree gaps formed; Mullineaux et al. 2016, Alvarez-Buylla and Garcia-Barrios 1993). But just as easily could capture the rates at which rock pools desiccate relative to the generation and colonization times of unicellular algae, bacteria, daphnia and copepods or fish (e.g. Altermatt et al. 2012, Bengtsson 1993). Overall, our simulations capture habitat dynamics representative of systems where all habitat is near constantly present to highly variable within the timescales on which ecological systems may be observed. Each of these sets of simulations were repeated for species varying in dispersal ability from an average dispersal capability across the entire landscape (global dispersal), to an average capability of reaching the next nearest location of habitat formation (stepping stone), to an average of 1/10th this stepping stone dispersal. The dynamic habitat landscape across simulations consisted of a linear array of equidistant locations at which patches could be formed or lost with homogenous carrying capacities. However, the model is also capable of simulating more realistic landscapes akin those used in Walker and Gilbert (2022), Hanski and Ovaskainen (2000), and Rubio et al. (2014).

All simulations were initialized in a habitat configuration chosen with probability given by the quasi-equilibrium distribution and with *p*_0_ = *p*^*^ within that initial habitat configuration. This means simulations begin at quasi-equilibrium removing any need for a burn in period. Using an approach akin to Walker and Gilbert (2022) to keep simulated metapopulation dynamics comparable across landscapes and disturbance regimes, local extinction rates were to set (by adjusting *e* to set *δ* = *e*/*c*)to achieve a common landscape capacity in the stochastic landscape 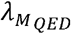to an equivalent value of 20 in every simulation. This ensured simulated metapopulations were expected to be similarly viable within landscapes differing in disturbance regime.

## Results

### Habitat distribution and the quasi-equilibrium distribution

The QED described the spatiotemporal distributions of the amount and configuration of habitat across the full parameter space of our simulations. This can be seen in Fig. 2 where the expected geometric mean number of habitable patches at QED is always well within the interquartile ranges of the geometric mean number of habitable patches throughout simulations. At one extreme when rates of disturbance relative to recovery are low, habitat is rarely lost and recovers extremely quickly (e.g. only a single patch was lost after ∼7000 “years” yet recovered within a “year” in Fig. S1A). This results in dynamic landscapes that are nearly static with all possible habitat fully present. When rates of habitat disturbance and recovery are more similar, landscapes become more variable and the amount of habitat may fluctuate on very short timescales the faster disturbance occurs relative to habitat recovery (Fig. S1B&C). Equal but faster rates of disturbance and recovery do not change the QED (Fig S1B versus C). For example, in the simulation selected for Fig. S1B, the absorbing state was reached shortly after 150 “years”. Since the QED is expected to describe the amount of habitat conditioned on some habitat existing to support a metapopulation, we can see the geometric mean amount of habitat matches or is very close to the simulated amount of habitat until it was all lost. At the furthest extreme, where habitat is lost quickly and recovers slowly, variation in the amount of habitat increases while also resulting in less habitat on average, which also causes it to be available for a shorter amount of time (Fig. S1D). Even at this extreme, however, average time until reaching a no-habitat state occurs on the order of 10,000 “years”.

**Figure 2.**
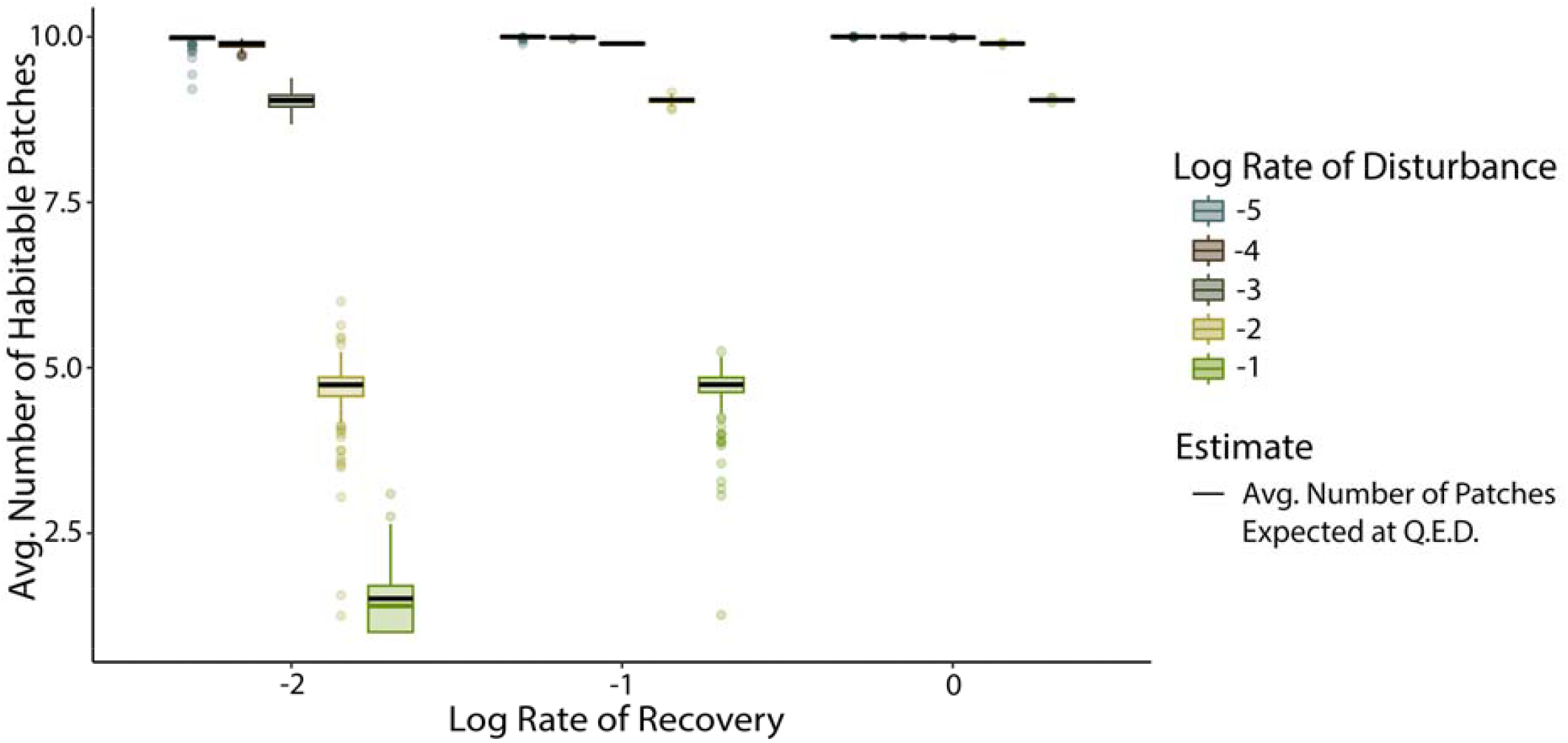
Expected versus simulated geometric mean amount of habitat across disturbance regimes varying in rate of habitat patch recovery and destruction. Black lines correspond to the expected amount of habitat size as weighted by the quasi-equilibrium distribution of habitat configurations and box plots display the median and 1.5x the interquartile range of the simulated amount of habitat (N = 200).

### Metapopulation size with habitat impermanence

Our simulations of metapopulations within impermanent landscapes show that the quasi-equilibrium occupancy, to *p*^*^_*QED*_, performed better than previous metrics across a range of impermanent landscapes and dispersal levels (Figs. 3-5). The closer match of *p*^*^_*QED*_to simulated metapopulation sizes, and the inferior performance of the implicit spatiotemporal averaging approach *p*^*^_*DEWoody*_ illustrate that it is necessary to account for the frequency and duration of spatial configurations the habitat may take through time. In the extreme cases where rates of habitat recovery greatly exceed rates of habitat loss, *p*^*^_*QED*_ performs similarly to *p*^*^_*DEWoody*_.

**Figure 3.**
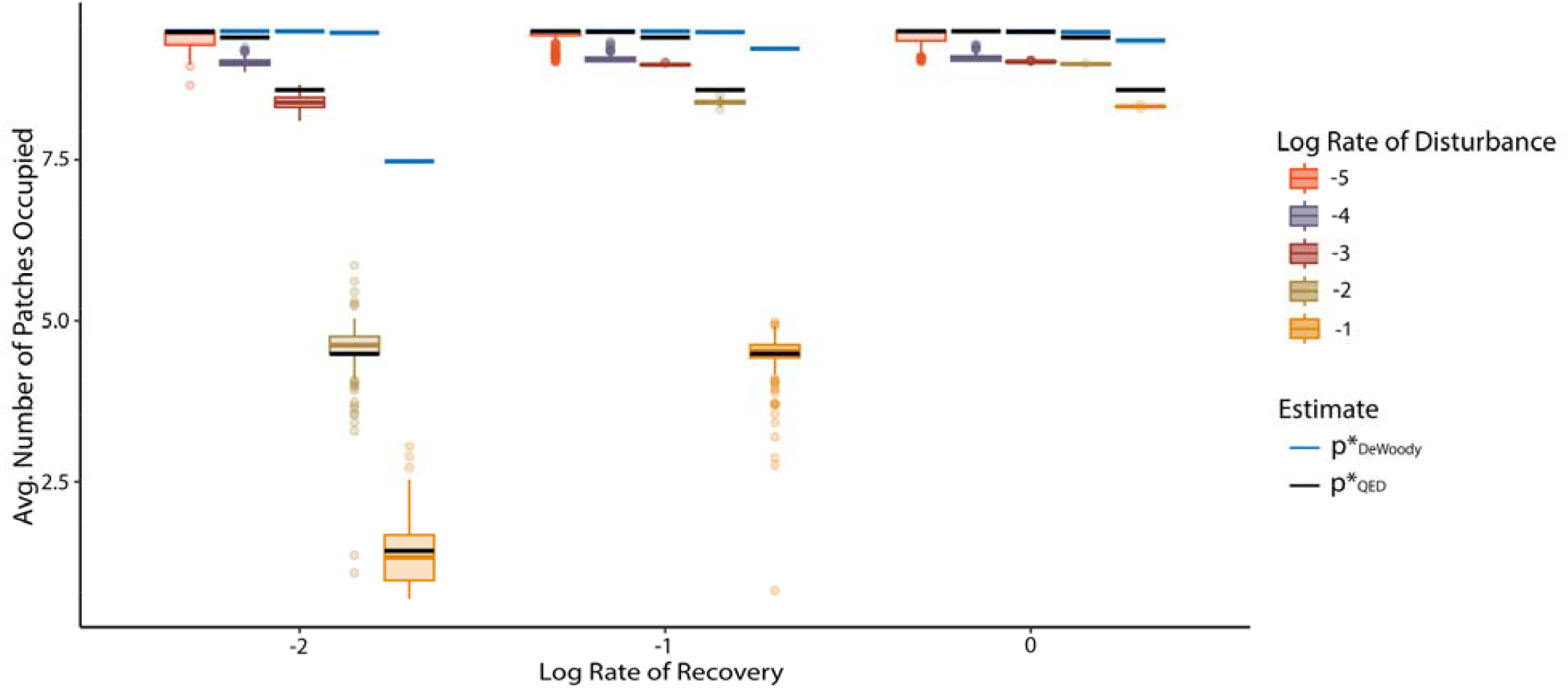
Metapopulation occupancy with habitat impermanence when individuals disperse across the whole landscape on average. Simulated (boxplots) and expected (black and blue lines) mean metapopulation occupancy across disturbance regimes varying in rate of habitat patch recovery and disturbance. Each data point is the average of a single simulation (N = 200 per habitat rate combination).

**Figure 4.**
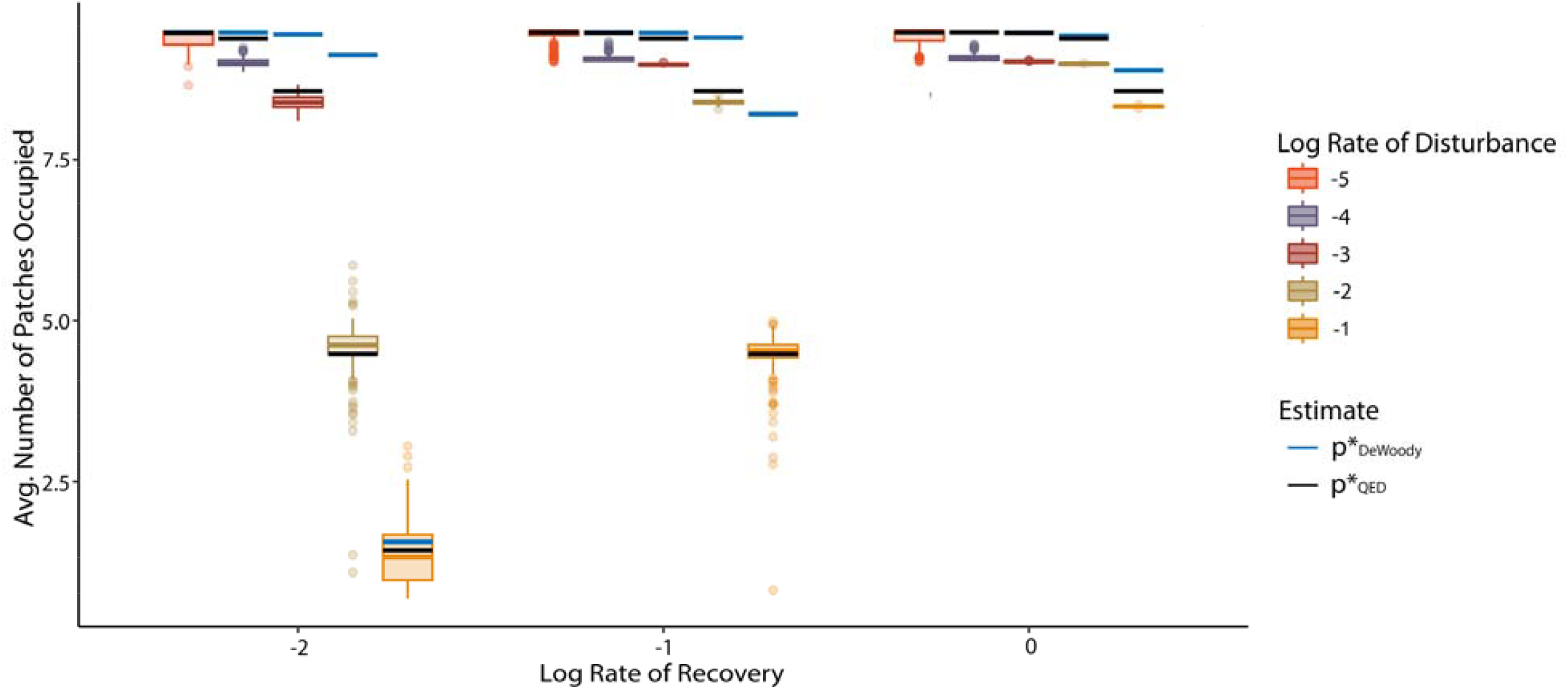
Metapopulation occupancy with habitat impermanence when mean dispersal distance is equal to the nearest potentially viable patch. Simulated (boxplots) and expected (black and blue lines) mean metapopulation occupancy across disturbance regimes varying in rate of habitat patch recovery and disturbance. Each data point is the average of a single simulation (N = 200 per habitat rate combination).

**Figure 5.**
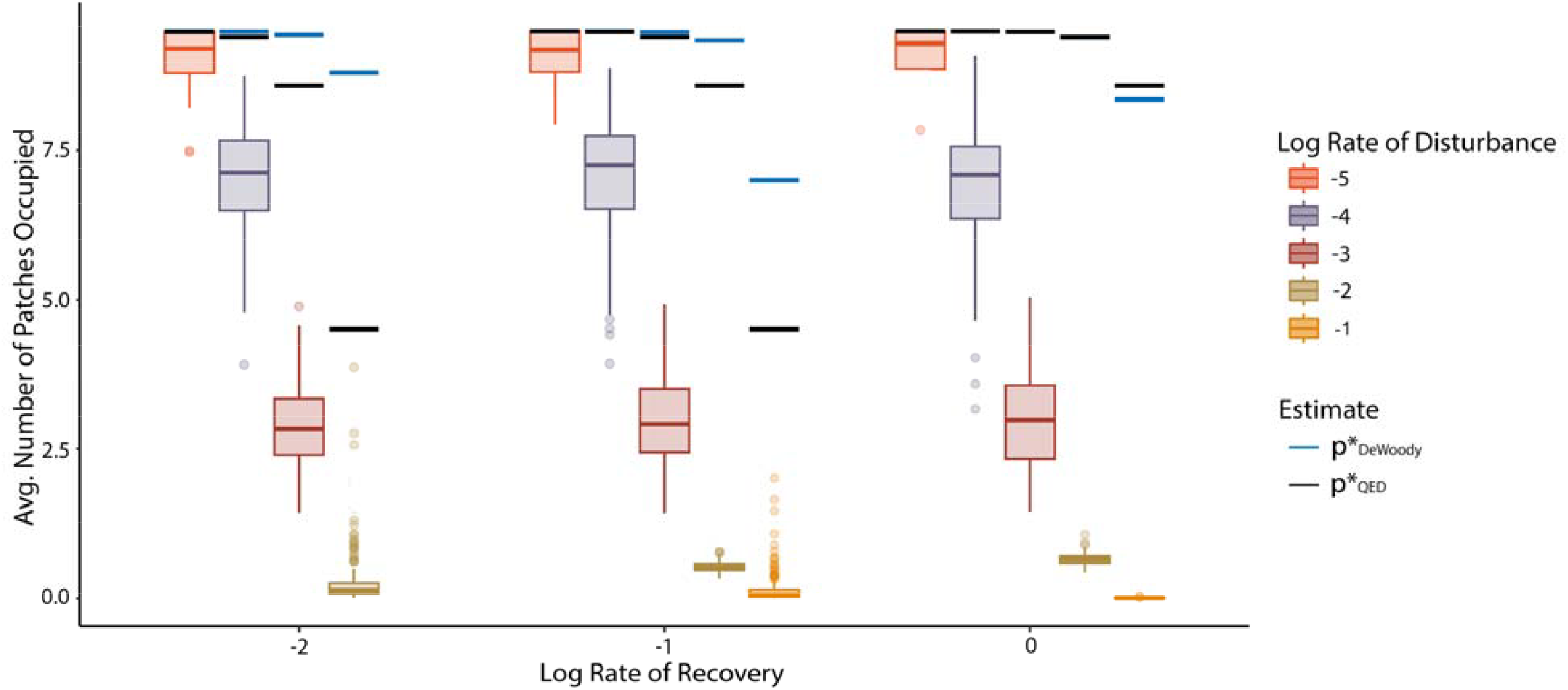
Metapopulation occupancy with habitat impermanence when mean dispersal distance is equal to 1/10^th^ the nearest potentially viable patch. Simulated (boxplots) and expected (black and blue lines) mean metapopulation occupancy across disturbance regimes varying in rate of habitat patch recovery and disturbance. Each data point is the average of a single simulation (N = 200 per habitat rate combination.

However, this equivalence only occurs when all possible habitat is present nearly all the time (e.g. fig. S1C). When metapopulation dispersal was not limited, meaning that species could disperse across the entire landscape, connectivity is altered the least by changes in habitat configuration. Thus, both metrics approach the *p*^*^ of a complete and static landscape (where all habitat patches are always habitable), in this case where nearly all the habitat is occupied at all times (fig. S2C). However, when habitat recovery rates are at all similar to disturbance rates (even still differing by an order of magnitude), impermanence significantly decreases metapopulation sizes (fig. S2B&C). This decreases average patch lifespans, decreasing *p*^*^_*DEWoody*_, but the spatiotemporal distribution of habitat configurations further reduces metapopulation sizes. Thus, our estimate provides closer predictions of average simulated occupancy, although its reliability is influenced by dispersal limitation and fast habitat dynamics.

Fast habitat dynamics and dispersal limitation caused the mean metapopulation size to deviate below what would be expected if transients were absent, as described by *p*^*^.

Fast habitat dynamics and dispersal limitation caused the mean metapopulation size to deviate below what would be expected if transients were absent, as described by *p*^*^_*QED*_. This can be seen even with global dispersal when habitat dynamics are fastest (Fig. 3 cases furthest to the right). When dispersal is more limited, such that populations disperse from one patch to the next possible patch on average (Fig. 2), or are rarely able to disperse to neighboring patches (Fig. 5), fast habitat dynamics cause an even more marked decrease in metapopulation sizes. In general, observed occupancy (p) varied less than *p*^*^within simulated habitat configurations suggesting transients overall limited metapopulations from reaching equilibria within habitat configurations (Table 1). Highly limited dispersal, which is only viable when extinction rates (*e*) are low, showed the opposite pattern (Table 2 asterixed cases), suggesting transients may more often prolong impacts of extreme configurations in these regimes. However, on average we see no evidence for any extended benefits of good configurations ever outweighing the costs of poorer configurations to produce higher average occupancies than predicted at QED (Figs. 3-5). When dispersal is extremely limited and habitat disturbance is much faster than habitat recovery this can cause metapopulation extinctions even when metapopulations are otherwise expected to persist at QED 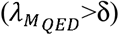 (e.g. Fig. 5 B&C).

**Table 1.** Mean time to absorption(T_*abs*_; no-habitat state) and mean variance of equilibrium occupancy (*p*^*^)and observed occupancy (*p*^*^) within simulations (N = 200 per habitat rate combination). Results are for a globally dispersive species in habitats differing in rates of disturbance (*r*_*off*_) and recovery(*r*_*on*_).

| $\log(r_{on})$ | $\log(r_{off})$ | $T_{abs}$ | Variance of $p^*$ | Variance of $p$ |
| --- | --- | --- | --- | --- |
| -2 | -5 | $>1 \times 10^{20}$ | 0.23 | 0.17 |
| -2 | -4 | $>1 \times 10^{20}$ | 0.35 | 0.30 |
| -2 | -3 | $1.5 \times 10^{13}$ | 0.98 | 0.75 |
| -2 | -2 | $1.2 \times 10^6$ | 2.49 | 2.00 |
| -2 | -1 | $1.1 \times 10^4$ | 0.48 | 0.27 |
| -1 | -5 | $>1 \times 10^{20}$ | 0.21 | 0.16 |
| -1 | -4 | $>1 \times 10^{20}$ | 0.27 | 0.24 |
| -1 | -3 | $>1 \times 10^{20}$ | 0.34 | 0.30 |
| -1 | -2 | $1.5 \times 10^{13}$ | 1.00 | 0.78 |
| -1 | -1 | $1.2 \times 10^6$ | 2.44 | 1.88 |
| 0 | -5 | $>1 \times 10^{20}$ | 0.21 | 0.16 |
| 0 | -4 | $>1 \times 10^{20}$ | 0.26 | 0.24 |
| 0 | -3 | $>1 \times 10^{20}$ | 0.26 | 0.24 |
| 0 | -2 | $>1 \times 10^{20}$ | 0.34 | 0.31 |
| 0 | -1 | $1.5 \times 10^{13}$ | 0.99 | 0.83 |

**Table 2.** Mean variance of equilibrium occupancy(*p*^*^)and observed occupancy(*p*) for a species with an average dispersal distance equal to the interpatch distances at which habitat is gained and lost (Stepping Stone), and a species with 1/10^th^ this dispersal distance on average (1/10^th^ Stepping Stone) in habitats differing in rates of disturbance(*r*_*off*_) and recovery(*r*_*on*_). (N = 200 per habitat rate combination). *’d values indicate when p varied more amongst simulations than *p*^*^.

| | | Stepping Stone | | $1/10^{\text{th}}$ Stepping Stone | |
| --- | --- | --- | --- | --- | --- |
| $\log(r_{on})$ | $\log(r_{off})$ | Variance of $p^*$ | Variance of $p$ | Variance of $p^*$ | Variance of $p$ |
| -2 | -5 | 0.23 | 0.17 | 0.21 | 0.06 |
| -2 | -4 | 0.35 | 0.30 | 0.33* | 1.35* |
| -2 | -3 | 0.98 | 0.75 | 0.96* | 3.62* |
| -2 | -2 | 2.49 | 2.00 | 2.45 | 0.28 |
| -2 | -1 | 0.48 | 0.27 | 0.33* | 0.64* |
| -1 | -5 | 0.21 | 0.16 | 0.20 | 0.10 |
| -1 | -4 | 0.27 | 0.24 | 0.27* | 1.74* |
| -1 | -3 | 0.34 | 0.30 | 0.34* | 3.80* |
| -1 | -2 | 1.00 | 0.78 | 0.99 | 0.50 |
| -1 | -1 | 2.44 | 1.88 | 2.44 | 1.88 |
| 0 | -5 | 0.21 | 0.16 | 0.19 | 0.10 |
| 0 | -4 | 0.27 | 0.24 | 0.27* | 1.74* |
| 0 | -3 | 0.26 | 0.24 | 0.26* | 3.68* |
| 0 | -2 | 0.34 | 0.31 | 0.33* | 0.64* |
| 0 | -1 | 0.99 | 0.83 | 0.99 | 0.02 |

## Discussion

Our study provides new insights on the complex impacts that habitat impermanence has on metapopulation dynamics, even in simple landscapes. Our results caution against using simple averages to estimate the impacts of habitat impermanence in spatially heterogenous landscapes – accounting for the spatiotemporal distribution of habitat invariably improves estimates of metapopulation size. However, while considering the full spatiotemporal connectivity of habitat improves metapopulation metrics, it is not sufficient to accurately describe dynamics for more dispersal limited species, especially when rates of habitat destruction are high. Importantly, non-equilibrium metapopulation dynamics produce outcomes that metapopulation estimates may often miss. The extent to which these complex impacts become important may instead be predicted through understanding how species traits modify the rates of metapopulation dynamics relative to rates of habitat change. Because species and habitat rates can differ, transient dynamics are likely to play a role in the size and persistence of many metapopulations. These insights highlight new concerns, but also potential benefits, of habitat impermanence for population management and conservation.

### Performance of Metapopulation Estimates in Impermanent Habitat

Our approach provides estimates of metapopulation viability and size by fully accounting for the spatiotemporal connectivity of habitat through impermanence of habitat patches – a goal that theory has increasingly called for and yet has also been criticized for failing to capture (DeWoody et al. 2005, Van Teeflelen et al. 2012, Perry and Lee 2019). It is likely that the poor performance of metrics that homogenize spatiotemporal distributions of habitat improve in much larger and well-connected habitat networks than we explored, as predicted for infinitely large metapopulations (McVinish et al. 2016). However, in finite metapopulations with even modestly limited dispersal, capturing the full spatiotemporal connectivity of habitat becomes necessary.

Indeed, even though fully accounting for spatiotemporal connectivity improves key metrics, our work also shows that these metrics are inaccurate and may be insufficient for conservation and management of many species.

Empirical tests of metapopulation dynamics that incorporate habitat impermanence are rare, but at least one study has taken an approach that is conceptually similar to our framework for spatiotemporal dynamics. Bertasello and colleagues tested the dependence of metapopulation size on the fluctuating availability of wetland habitats for several species by first using hydrodynamics to assess the observed spatiotemporal distribution of available habitat, and then calculating expected occupancies weighted by the available habitat (Bertassello et al. 2021).

While their underlying model differed in its complexity and averaging approach, it similarly overestimated metapopulation sizes. Given that a fully spatiotemporal approach tends to predict more conservative occupancies than that of previous authors (Figs. 3-5), it appears that a full spatiotemporal accounting of habitat is better, but imperfect, in empirical settings. Caution must be taken even when full characterization of the spatiotemporal distribution of habitat configurations is possible because equilibria solutions fail to incorporate the effect of transients on metapopulation size and viability.

In theory, transient dynamics can exacerbate impacts of local extirpations with patch loss, offset them by slowing losses from remaining patches, or even cause fluctuations in occupancy in some circumstances (Gurney and Nisbet 1978). For the spatially realistic Levins model, the rates of colonization and extinction determine transients by setting the speed at which the metapopulation responds to new habitat configurations. Intuitively, decreased dispersal slows metapopulation dynamics via its effect on colonization; when this slowing causes the speed of habitat changes to outpace metapopulation growth, the metapopulation will spend less time at the predicted equilibria and may also fail to reach equilibria altogether. Since extirpation with patch loss is immediate while increased colonization can only offer decreased delay of metapopulation gain, increased transients through slowed colonization intuitively bias metapopulation sizes lower. Conversely, to achieve the same level of viability (expected 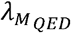) in landscapes that hold fewer habitat patches on average, metapopulations must have lower local extinction rates, which slow metapopulation dynamics through the speed of non-habitat caused local extinction. This slows the rate at which occupied patches empty when habitat configurations become poorer, a phenomenon that has been shown to cause metapopulations to persist long past the time when they are expected to go extinct following habitat destruction (Tilman et al. 1994; Hanski and Ovaskainen 2002). Nonetheless, our results overall show these possible benefits of decreased local extinction rates fail to counter the increased habitat-induced extinctions that occur at high patch loss and recovery rates.

Transients may always have this negative impact on metapopulations since, in addition to rates of colonization and extinction, the proportion of occupied habitat (p) can feedback to influence transient dynamics. In a fully connected metapopulation, the maximum rate of change in the metapopulation occupancy occurs when p is at half its equilibrium value, suggesting that large perturbations in p (away from p*) could induce slower dynamics in some landscapes. This effect is similar to that seen in populations experiencing logistic growth (Law et al. 2003). Although the rate becomes more complex when spatial heterogeneity limits dispersal amongst patches (Law et al. 2003), examination of eqn. 10 shows the multiplicative importance of nearby patches on the metapopulation rate of change that underlies the effect of p on colonization.

## Broader Ecological Implications

We provide this means of accurately characterizing spatiotemporal connectivity in hopes such methods become more widely considered and adopted in the conservation and management of species residing within impermanent habitats. Metapopulation viability and connectivity metrics have become commonplace in management and have informed standard techniques for empirically estimating importance of habitat connectivity and quality to species (DeAngelis and Yurek 2017, Saura and Rubio 2010). However, studies risk underestimating the importance of habitat connectivity and quality to these species when they fail to consider habitat impermanence (Hodgson et al. 2009). Meanwhile, incorporating spatial connectivity of landscapes with the average temporal habitat availability has led to potentially misleading conclusions that habitat impermanence necessarily impacts metapopulations primarily through patch lifespans (Ross et al. 2008, McVinnish et al. 2016, Van Teeffelen et al. 2012). The common logical extension of this conclusion is that metapopulations may be better maintained through efforts focused on patch preservation rather than improving spatial connectivity. Our research shows that this is misleading by illustrating how spatial and temporal connectivity jointly influence metapopulations. Despite the more detailed and accurate assessment achieved with our methods, they require the same information on landscapes as previous, homogenizing methods: where and how frequently habitat is lost and formed in landscapes, and population estimates common to all metapopulation models. Even knowledge of a subset of the most frequent configurations in landscapes could offer the opportunity to dramatically improve estimates of metapopulation viability and size (see Appendix).

We see here that understanding species traits relative to rates of change in their landscapes will inform us of the species and conditions under which transients are, or become, important. Dispersal ability is hard to quantify, yet critical, and must be quantified relative to the distribution of habitat (Rayfield et al. 2023, Walker and Gilbert 2022). However, understanding even the degree to which species may be capable of reaching locations of potentially viable habitat may be informative. Likewise, we see life history traits that impact rates of colonization and extinction are critical, and are most relevant when considered relative to rates of habitat change.

Because transients are generally detrimental, our work agrees with previous conclusions that preserving patch lifespans should greatly benefit individual species (Ross et al. 2008, Van Teeffelen et al. 2012). However, our results provide more nuance by demonstrating that where preservation efforts are focused in a landscape is important, may even shift through time, and may depend on whether a patch currently supports a local population. Some earlier studies have concluded that increasing number of impermanent habitat patches offers little benefit to metapopulations due to the disproportionate effect of patch lifespan on viability and size metrics (Van Teeffelen et al. 2012). However, strategic placement of impermanent habitats could still be most cost effective and key to preserving some metapopulations by allowing colonization rates to better ‘keep up’ with fast habitat dynamics (Figs. 3 versus 5 when more habitat is available on average). Adopting metrics that quantify spatiotemporal connectivity will be key to choosing appropriate land management strategies based on different feasibility constraints particular to different ecological systems.

Our study also suggests possible benefits of habitat impermanence in multispecies communities that deserve further study. As detrimental as impermanence of one habitat type may be for a single species, it may be critical to the persistence of another (e.g. for early versus late successional species) (Amarasekare and Possingham 2001). Furthermore, because we see that transients may limit metapopulation growth rates, they may impose additional consequences on coexistence of species within even the same habitats. These dynamics would be especially beneficial for weaker competitors if a competition-colonization trade-off aids coexistence (Tilman 1994). The nonlinearity of the dynamics underlying metapopulation size suggests that prolonging habitat states could cause a metacommunity to cross a critical threshold at which a species shifts from relatively low occupancies to dominating entire landscapes.

## Approach and Model Limitations

A major limitation to our results is that our study was done only for small, finite landscapes. Meanwhile approaches homogenizing spatiotemporal dynamics hold in infinite landscapes (McVinnish et al. 2016). This leaves the effect of spatiotemporal habitat dynamics in larger but finite landscapes unknown and worthy of further investigation. A major barrier to our approach is the exponential increase in computational time and memory required to obtain the QED of habitat configurations when increasing the number of feasible patches within a landscape. In the case where patch loss and recovery is random and independent, the number of habitat states tracked by the CTMC can be reduced from 2^*N*^ -1 to N since each habitat configurations with an equal number of patches available have the same probability at QED. Our metrics, 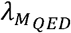 and *p*^*^*QED*, can then be estimated using a subset of the most frequent configurations at QED. Indeed, further exploration of the sensitivity to the fraction of possible configurations used suggests ∼20% of configurations may be sufficient to recover 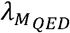 and *p*^*^_*QED*_ based on the full set of configurations (Appendix Fig. S5). However, computational constraints remain for simulating and recording metapopulation dynamics in large landscapes. Increasing computational advances and resources are expected to decrease these challenges in the near future. Thus, obvious future directions for this work include exploring the sensitivity of our results to networks with greater numbers of patches.

Another major limitation of our study is that it was done on minimally complex landscapes unlike those found in nature. This was intentional for the sake of providing clear evidence for the importance the spatiotemporal distribution of habitat in finite landscapes even when potential habitat quality, sites and rates of gain and loss are all homogenous. However, many landscapes show highly unequal patch contributions in supporting a metapopulation through clumped distribution of habitat of variable quality (Hanski and Ovaskainen 2000). Habitat impermanence creates spatially heterogenous landscapes even when locations of possible habitat gain and loss are distributed evenly (Fig. 1), and may well have a larger effect in already heterogeneous landscapes. However, it is not obvious how transients, especially if patch dynamics were correlated, may in turn respond to this increased heterogeneity.

## Conclusion

Our work overall suggests that spatiotemporal connectivity of habitat deserves greater attention as an important feature of landscapes, and that metapopulation research must better capture the interactive effects of temporal and spatial heterogeneity. Our approach provides improved metrics to assess metapopulation size and viability within impermanent habitats, but also highlights the importance of transient dynamics and the need to better understand and predict their impacts. Meanwhile, empirical work needs to quantify extinction and colonization rates relative to rates of habitat loss and gain in order to better understand extant dynamics and predict the consequences of changing rates in disturbed habitats.

## Appendix

**Figure S1.**
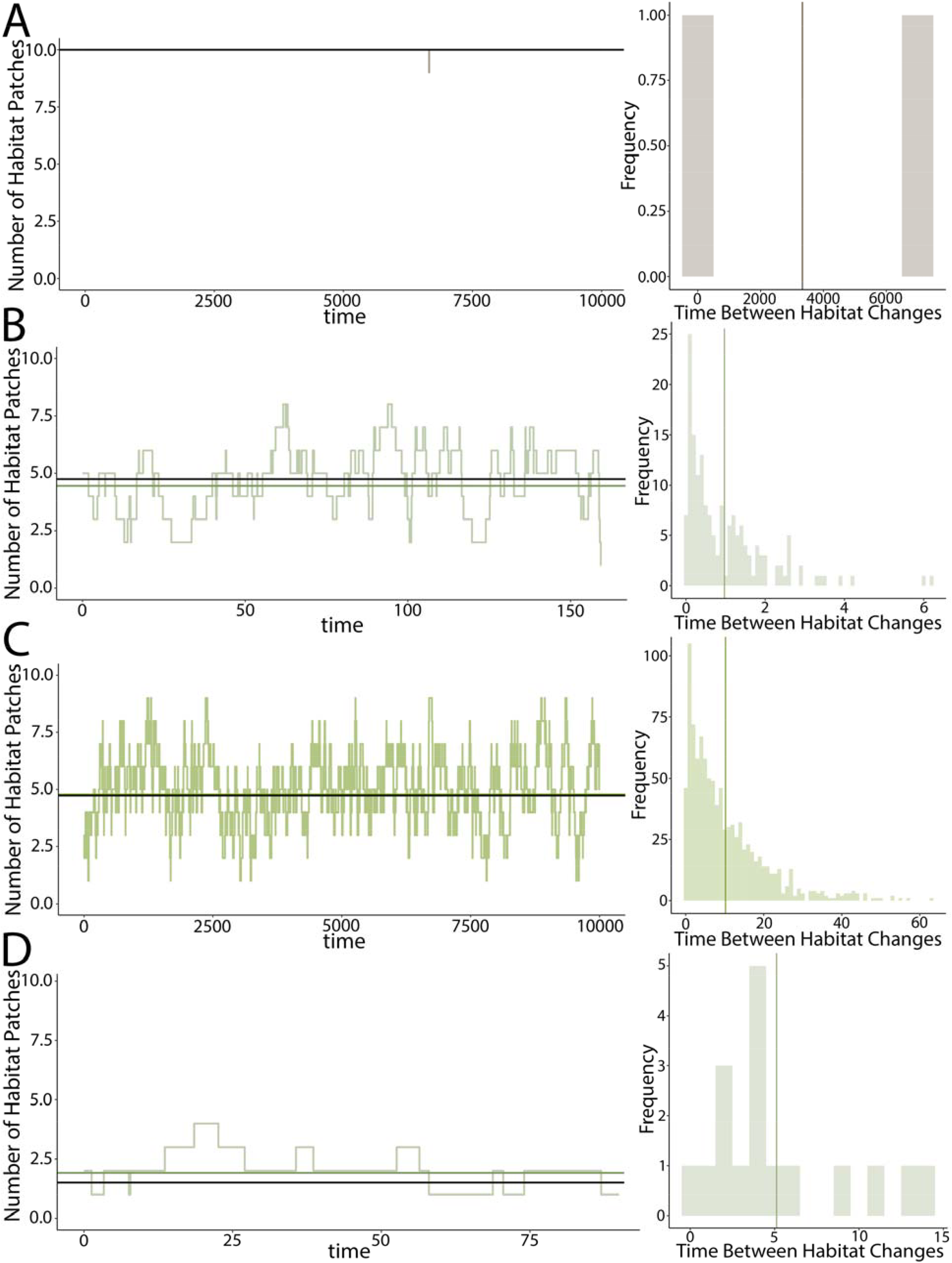
Examples of habitat dynamics within simulations providing the amount of habitat through time and a histogram of the distribution times between habitat changes (*τ*) (lighter shades) and their average (darker lines) when A) *r*_*off*_ = 1*x*10^−5^ and *r*_*on*_ = 1, B) *r*^*off*^ = 1*x*10^−2^ and *r*_*on*_ = 1*x*10-2, C) *r*_*off*_ = 1*x*10^−1^ and *r*_*on*_ = 1*x*10^−2^, and D) r*off* = 1*x*10-1 and *r*_*on*_ = 1*x*10-1.

**Figure S2.**
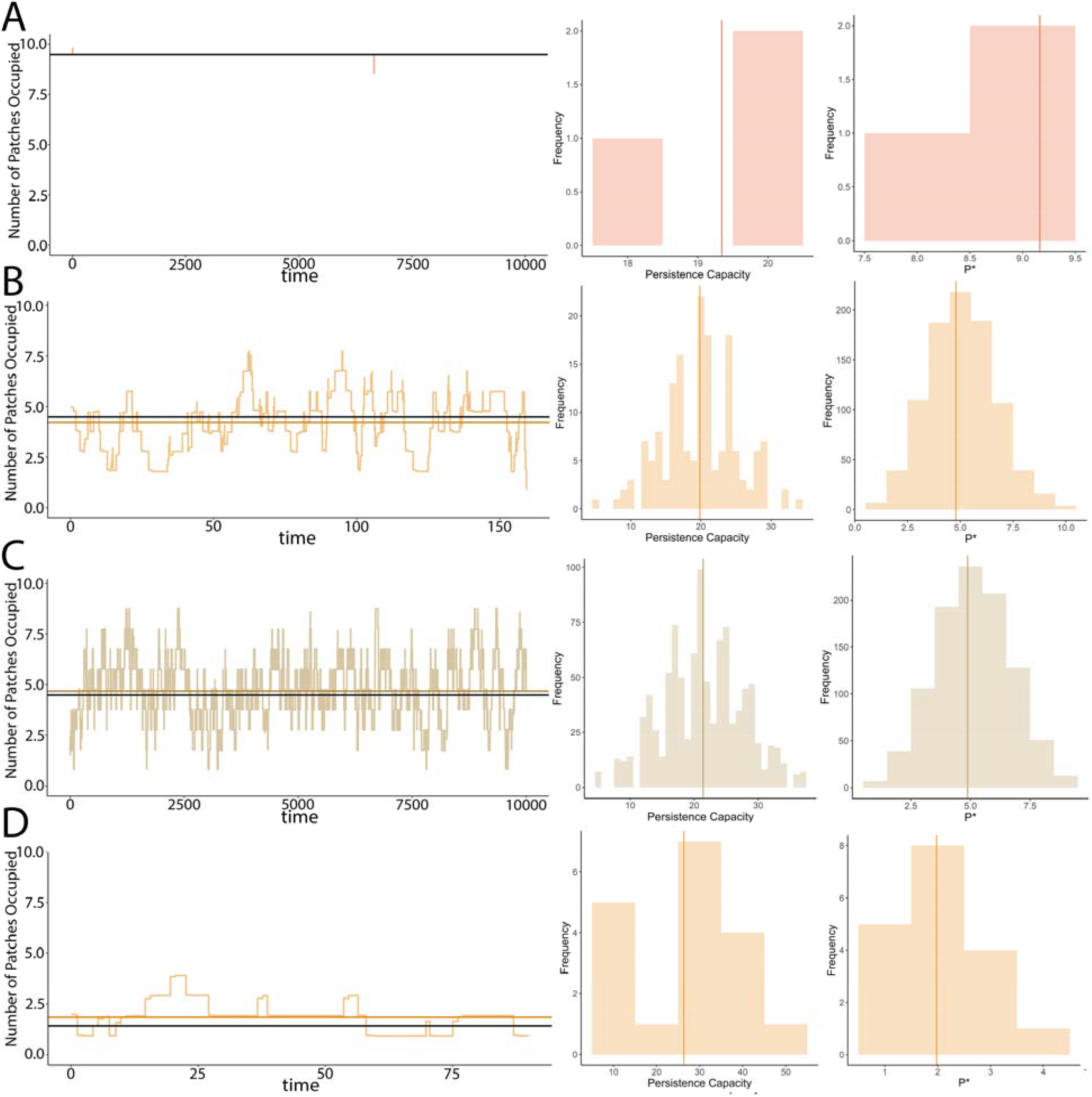
Examples of metapopulation dynamics with habitat impermanence within selected simulations when individuals disperse across the whole landscape on average, with (occupancy, *p*) through time, darker horizontal lines indicating its average and black lines indicating 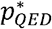. Histograms of the λ_*M*_’s and *p*^*^’s within simulated configurations with their averages shown by darker lines when A) *r*_*off*_ = 1*x*10^−5^ and *r*_*on*_ = 1, B) *r*_*off*_ = 1*x*10^−1^ and *r*_*on*_ = 1*x*10^−2^, C) *r*_*off*_ = 1*x*10^−1^ and *r*_*on*_ = 1*x*10^−2^, and D) *r*_*off*_ = 1*x*10^−1^ and *r*_*on*_ = 1*x*10^−1^.

**Figure S3.**
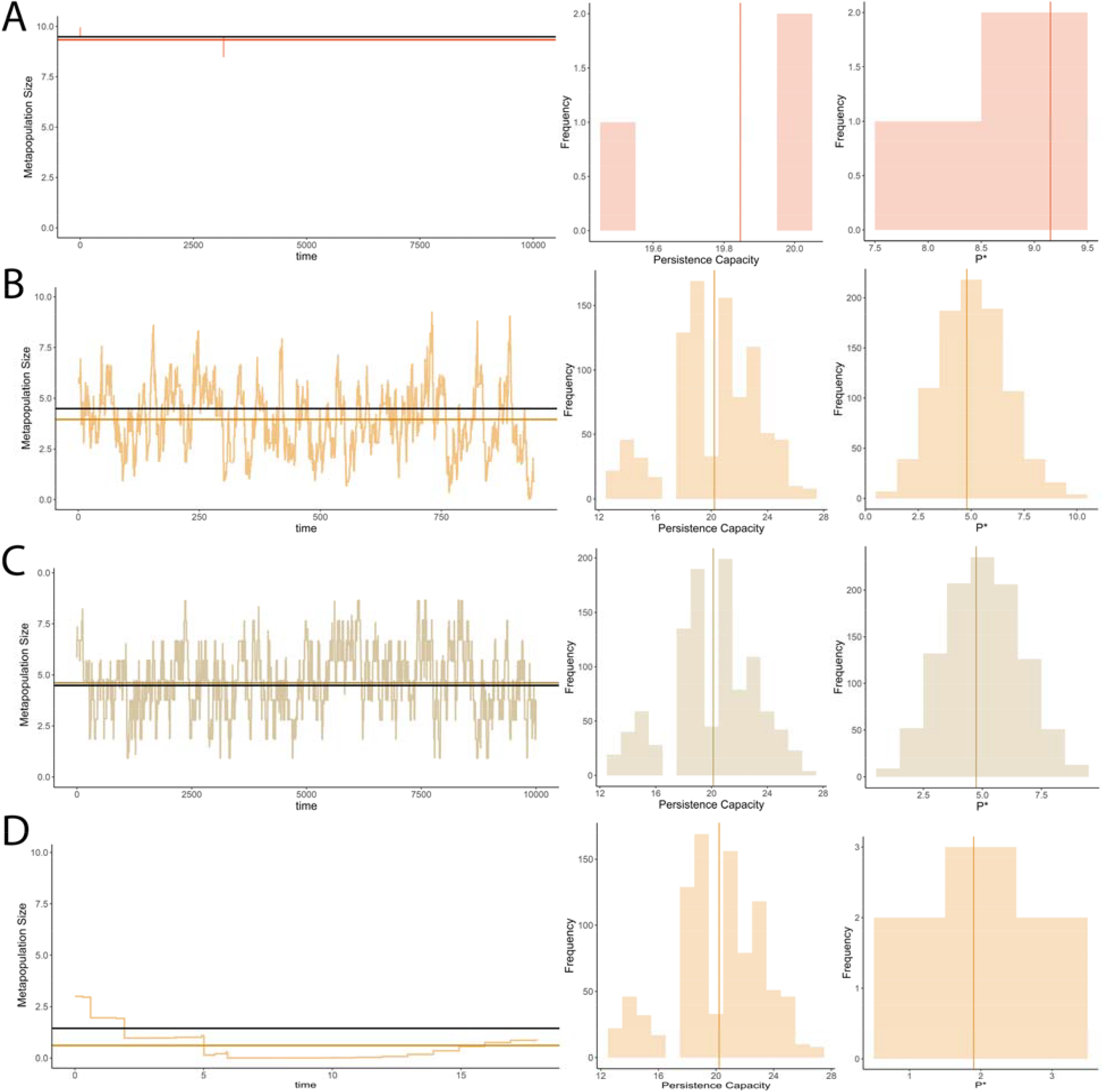
Examples of metapopulation dynamics with habitat impermanence within selected simulations when mean dispersal distance is equal to the nearest potentially viable patch, with (occupancy, *p*) through time, darker horizontal lines indicating its average and black lines indicating 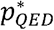. Histograms of the λ_*M*_’s and *p*^*^’s within simulated configurations with their averages shown by darker lines when A) *r*_*off*_ = 1*x*10^−5^ and *r*_*on*_ = 1, B) *r*_*off*_ = 1*x*10^−2^ and *r*_*on*_ = 1*x*10^−2^, C) *r*_*off*_ = 1*x*10^−1^ and *r*_*on*_ = 1*x*10^−2^, and D) *r*_*off*_ = 1*x*10^−1^ and *r*_*on*_ = 1*x*10^−1^.

**Figure S4.**
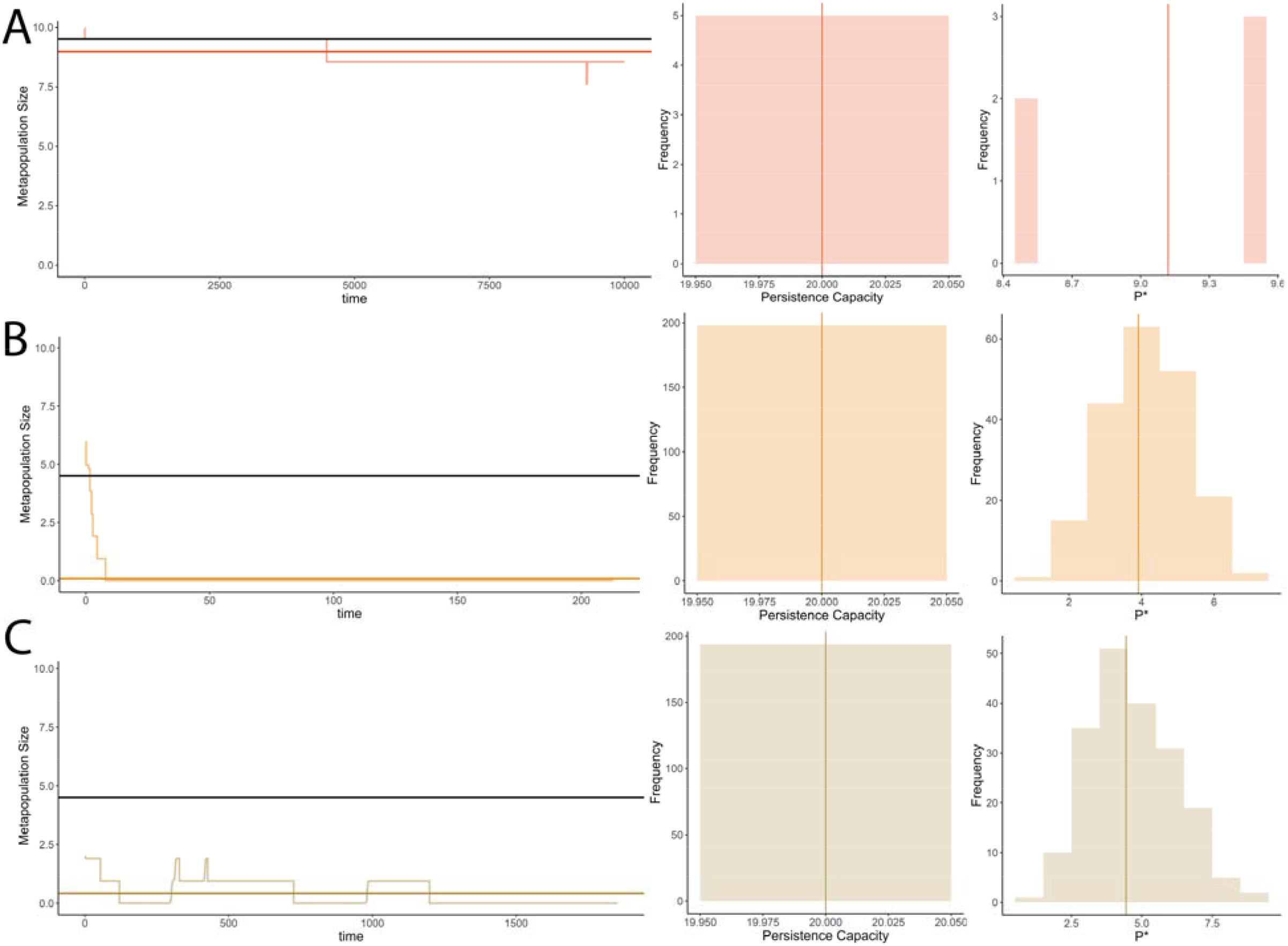
Examples of metapopulation dynamics with habitat impermanence within selected simulations when mean dispersal distance is equal to 1/10^th^ the nearest potentially viable patch, with (occupancy, *p*) through time, darker horizontal lines indicating its average and black lines indicating 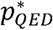. Histograms of the λ_*M*_’s and *p*^*^’s within simulated configurations with their averages shown by darker lines when A) *r*_*off*_ = 1*x*10 ^−5^ and *r*_*on*_ = 1, B) *r*_*off*_ = 1*x*10 ^−2^ and *r*_*on*_ = 1*x*10 ^−2^ C) *r*_*off*_ = 1*x*10^−1^ and *r*_*on*_ = 1*x*10^−2^, and D) *r*_*off*_ = 1*x*10^−1^ and *r*_*on*_ = 1*x*10^−1^.

**Figure S5.**
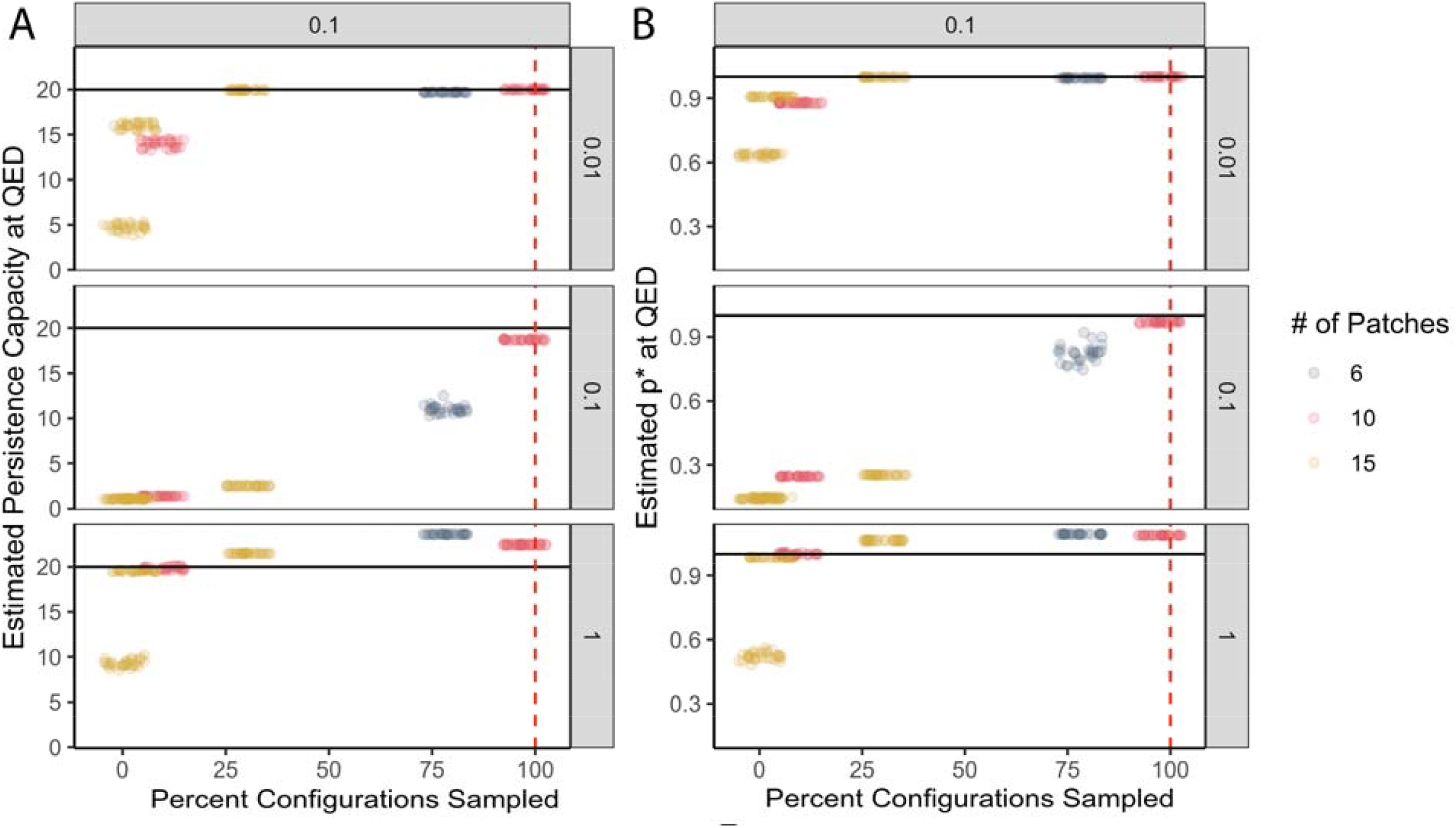
Sensitivity of estimating A) the persistence capacity at QED 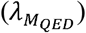 and B) the quasi-equilibrium occupancy 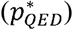 using a subset (percentage) of all possible configurations subsampled with probability of their occurrence at QED for a given rate of disturbance (*r*_*off*_, values displayed in top grey boxes) and recovery (*r*_*on*_, values displayed in right grey boxes). Data points have been horizontally jittered to better show replicate values using samples of 50 of all 64 possible configurations of 6 patches (78%), 100 (10%) and 1000 (98%) of 1024 possible for 10 patches, and 100 (0.3%), 1000 (3%), and 10000 (30%) of 32768 possible for 15 patches (N = 10). The horizontal black line indicates 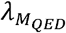’s true value as calculated using the full set of all possible configurations. The vertical dashed red line indicates where 100% of all possible habitat configurations would be sampled.

